# QuickFigures: a tool to quickly transform microscope images into quality figures

**DOI:** 10.1101/2020.09.24.311282

**Authors:** Gregory Mazo

## Abstract

Publications involving fluorescent microscopy generally contain many panels with split channels, merged images, scale bars and label text. Assembling and editing these figures with even spacing, consistent font, text position, accurate scale bars and other features can be tedious and time consuming. In order to save time and streamline the process I have created a toolset called QuickFigures.

## Introduction, Results, Discussion

Microscopy intensive scientific manuscripts often include figures with large numbers of fluorescent microscope images often split by channel into separate panels. A major publication can contain over 200 image panels! (1) To create a professional looking publication one must have consistent format within and between figures. However, commercial softwares do not provide quick ways to split channels and organize panels to generate figures without scores of error prone and repetitive mouse drags. Furthermore, major edits to figures like these are troublesome whether using Adobe Illustrator or Photoshop. For example, cropping or reordering over a dozen identically sized panels in Photoshop without mistakes or accidental changes in panel spacing cannot be done conveniently nor quickly. Although some tools for creating and formatting scientific figures have been created (2,3), we found that these did not suit our needs because they did not save time on many irksome but important steps. Therefore, I created QuickFigures, a set of tools that 1) can produce figures of sufficient quality for any journal, 2) is user friendly and easy to learn, 3) can save hours of time, 4) can export files into popular softwares like Adobe Illustrator, Microsoft Powerpoint, and Inkscape 5) is versatile enough to generate any conceivable style, layout, variation or format of figure, 6) is used as a free PlugIn for ImageJ, making adoption easy for researchers already familiar with ImageJ.

I have also created complete video tutorials (See Supplemental Materials, and <u>link</u>) as a guide to help first time users. This series of videos also clearly demonstrates the usefulness of QuickFigures. Below is a description of the figure production process.

After installing QuickFigures into ImageJ, new toolbars appear (**Figure 1**, see red arrows). Users can create a figure simply by first opening a multidimensional image stack in ImageJ, and then clicking the “Quick Figure” button that is visible on the Object Tools toolbar of Quickfigures (**Figure 1**, see purple arrow). This one click creates a figure with a series of split channel images, merged image, channel labels and scale bar. The user can edit every part of this figure. QuickFigures automates and facilitates mundane tasks that arise naturally during the editing process. Changes in panel spacing or panel order can be quickly applied to figures with dozens of panels. Insets can be created and edited easily with the Inset Tool (**Figure 1**, see Orange arrow and Movie S1). When one resizes an image panel, scale bars appropriately adjust. Text signifying each channel is colored and aligned automatically. Image resolution can be changed as needed. When a user adjusts the Min/Max of one channel, QuickFigures alters every image panel that contains that channel (Merge and Split) ensuring consistency of all adjustments across groups of multichannel images. Importantly, a user can set a default or “template” for figures to ensure that a single consistent format is automatically applied to newly created figures (Templates can also be applied to existing figures). In aggregate, several slow, or complex tasks are made fast and simple.

**Figure 1:**
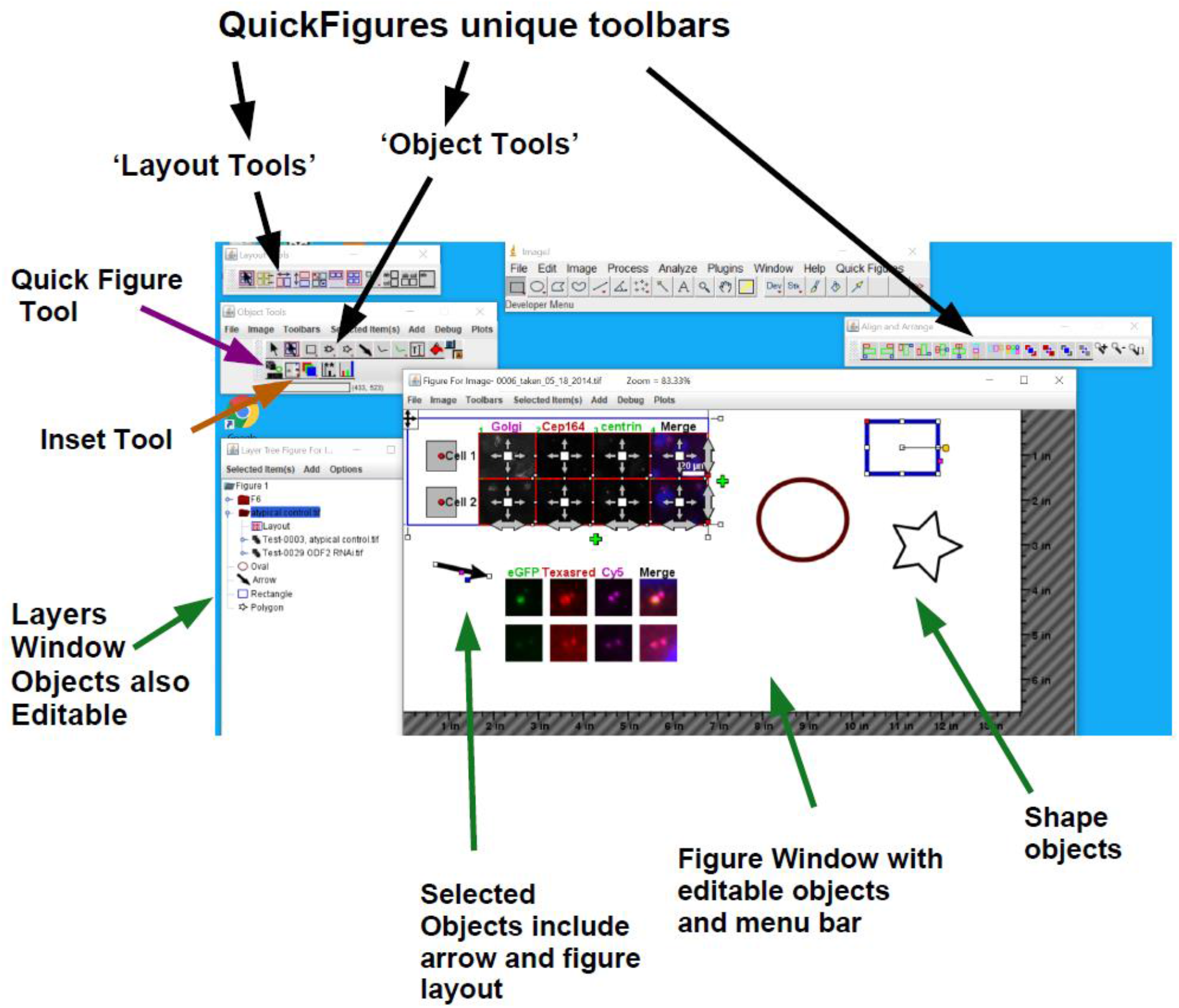
Appearance of ImageJ with QuickFigures installed. Key features of QuickFigures user interface are shown including the Figure, Toolbars and Layers Window.

QuickFigures was designed to feel familiar and logical to anyone who has used Powerpoint, Photoshop, Illustrator or any commercial software. Every item in QuickFigures is an editable object that can be clicked on, moved, resized, rotated, hidden, deleted, aligned, edited or duplicated (**Figure 1**, see green arrows). A simplistic set of menus and toolbars allows the user to add an unlimited number of items such as text, images, drawings, arrows, layers, plots, shapes and additional multichannel images. Right clicking and double clicking on items reveals popup menus and options dialogs for the clicked item. With minimal instruction, any researcher can understand the software and feel comfortable using it. The specialized features of QuickFigures are designed to require mere minutes explanation while saving time and providing convenience. After a user has created work in QuickFigures, a single click can generate the same figure in Adobe Illustrator. QuickFigures can also export files into Powerpoint (.ppt) or Scaleable vector graphic (.svg), a format that can be opened by many popular softwares including Adobe Illustrator. Exported figures are then suitable for further editing, presentation or publication.

## Materials and Methods

QuickFigures was written in Java, the same language as ImageJ. Existing ImageJ code was used for several key processes including 1) keeping track of an images spatial scale and units, 2) the assembly of channels into a merged image based on channel colors and display range, 3) scaling and rotating images (Using the bilinear interpolation algorithm), 4) keeping track of channel colors, channel order and metadata about channel names, exposure times 5) Serializing/saving ImageJ’s multichannel images. Since several complex processes could be performed using ImageJ, this eliminated the need to devise and implement new algorithms. However, the connection between QuickFigures core components and ImageJ was designed with an interface-based architecture such that ImageJ could be replaced by another package if future needs demand it. The user interface for QuickFigures was written using java foundation classes and can function on any operating system (Tested on Window and MacOsX). The QuickFigures package can also be reused to construct other softwares.

### Import and Export

The best (and most popular) tool for importing proprietary microscopy file formats into ImageJ is the **Bio-Formats Importer** created by the OME consortium (4) (See Movie S5 for use instructions and demonstration). This Importer also reads important metadata regarding spatial scale and channel names/channel colors. Assuming that a .Zvi, .Czi, .Lif or other microscopy format file is opened using Bio-Formats, QuickFigures will use the metadata as a basis for the initial channel labels within a figure. In order to test the accuracy of channel names, a series of example images stained with known markers in known colors was opened; QuickFigures’ channel labels were consistent with the channel names given by Bio-Formats. In order to export figures to ‘.ppt’ and ‘.svg’ file formats, two toolkits created by the Apache Foundation were used as libraries (POI for Powerpoint, https://poi.apache.org/ and Batik for SVG, https://xmlgraphics.apache.org/batik/). Instructions on how to install these libraries are included in the tutorial videos (Movie S5). To test the export, I created example figures with all types of objects including Images, Text, Scale Bars, multiple colors, arrows and other shapes. Exported figures resembled originals with matching details such as PPI of image panels, fonts, colors, width of lines etc.

### Other Testing

QuickFigures includes scores of objects, windows, menus, and components. Although most features could be tested simply by using them on multiple example figures, a few functions of QuickFigures demanded methodical testing. Of these, accuracy of scale bars was the most crucial feature. Since ImageJ maintains information on the spatial scale of images, that information is used by QuickFigures’s Image panel objects and scale bar objects. To test the accuracy, appropriate scale bar sizes for several images were first calculated manually and then compared to the sizes of QuickFigures scale bar objects. Subsequently, ImagePanels were resized to make sure that the scale bar objects appropriately changed size. Next, the units and spatial scale of the ImageJ Image was changed to make sure that the scale bar changed appropriately. In order to test complex features like the figure format menu commands, every possible type of figure edit was performed on a single figure and that figure’s format was saved. That saved format was then applied to a series of figures to confirm that the details of a saved figure format was indeed reflected in the target figure.

### Practical Use Test

After creation of an early version, QuickFigures was used constantly for preparation of figures for presentations for a period 3-5 years. Every format, layout and style of figure was generated for various scientific projects. During that time any irregularities, error messages and limitations that appeared were also fixed. Because of this practical testing, one can be confident that QuickFigures is suitable for widespread use.

### Installation

QuickFigures may be downloaded from <u>here</u>. QuickFigures can be installed into ImageJ simply by placing file into plugins folder of ImageJ. (see Movie S5)

## Supporting information

QuickFigures Software

## Acknowledgements

We acknowledge Professor Bryan Tsou. Images used for the demonstration videos and testing were taken using the microscopes within the Tsou Lab. We also acknowledge Dr. Brian O’Rourke who provided advice regarding this manuscript.

